# Incomplete letter recognition is limited by cortical and not optical factors: Simulating the visual deficits of dementia in healthy adults

**DOI:** 10.64898/2026.05.18.725904

**Authors:** Zien Huang, Tessa M. Dekker, Sebastian J. Crutch, Keir X. X. Yong, John A. Greenwood

## Abstract

Incomplete letter recognition tasks are frequently used to detect visual deficits arising from neurodegenerative syndromes, including Posterior Cortical Atrophy (PCA; ‘visual-variant Alzheimer’s disease’). A recent development of this approach is the Graded Incomplete Letters Test (GILT), which measures recognition thresholds for letters degraded by removing pixelated sections (decreasing ‘completeness’). Although GILT thresholds are strongly elevated in PCA relative to typical adults, the precise cortical visual impairments underlying these deficits are unclear, as is the potential contribution from age-related optical limitations. We compared candidate cortical factors (crowding and global integration) with optical limitations (blur and low contrast) by simulating these factors in typical adults (n=6) viewing incomplete letter stimuli. Participants identified foveally presented letters (12 alternatives), with completeness varied using QUEST. At baseline, thresholds averaged ∼5% completeness. Optical factors were simulated by separately applying blur and lowered contrast. These factors had minimal effect on thresholds, except where blur/contrast levels approached visibility limits, where thresholds rose modestly but remained far below clinical levels in PCA. Cortical factors were simulated by increasing crowding (disruptions from clutter) through peripheral presentation, with global-integration impairments simulated by varying pixel size to alter the distribution of degradation (limiting spatial integration) or degrading letters dynamically with limited-lifetime pixels (limiting temporal integration). These manipulations substantially elevated thresholds, with combined crowding and global-integration impairments increasing thresholds to levels comparable with PCA. We conclude that impaired incomplete letter recognition is driven primarily by cortical rather than optical factors, and that neurodegenerative deficits may reflect the combined impact of multiple cortical limitations.

## Introduction

Visual symptoms are a common feature of many neurodegenerative conditions (Armstrong et al., 2013; Blenkinsop et al., 2020; Jackson & Owsley, 2003; Tiraboschi et al., 2006), particularly in visual-led dementias such as Posterior Cortical Atrophy (PCA). PCA is a neurodegenerative syndrome characterised by impairments in everyday visual tasks including object recognition, reading, visual navigation, and visuo-motor integration (Crutch et al., 2012; Crutch et al., 2017; Yong et al., 2023), as well as acalculia (difficulty with numbers), alexia (difficulty reading), and anomia (difficulty retrieving object names; Tang-Wai et al., 2004). These difficulties are associated with deficits in visual brain areas of the parietal and occipital lobes, rather than the medial temporal areas typically associated with memory-led Alzheimer’s disease (AD; Crutch et al., 2012; Crutch et al., 2017). The visual symptoms of PCA are frequently misattributed to ocular or psychological conditions (Yong et al., 2023), often prompting repeated and unnecessary eyecare assessments (Crutch et al., 2012; Harding et al., 2018). Formal diagnosis is substantially delayed as a result, with an average of almost 4 years between symptom onset and diagnosis (Chapleau et al., 2024; van Vliet et al., 2012), limiting opportunities for early intervention and patient support. Accordingly, there is a clear need to improve the early detection of PCA via more sensitive and targeted diagnostic tests, which in turn requires a better understanding of the perceptual mechanisms underlying these visual impairments.

In both clinical and research settings, visual deficits are frequently measured using the recognition of visually degraded and/or ambiguous letters or objects, commonly in dementia and related neurodegenerative syndromes (Humphreys & Riddoch, 1993; Mioshi et al., 2006; Torfs et al., 2014; Warrington & James, 1991), as well as the diagnosis of PCA (Schott & Crutch, 2019; Yong et al., 2022). A widely used diagnostic tool for assessing these visual deficits is the Visual Object and Space Perception (VOSP) battery, a key part of which is the Incomplete Letters subtest, where letters are degraded by removing pixelated sections. The clinical utility of this subtest has led to its incorporation into widely used cognitive screening instruments for dementia, including the Addenbrooke’s Cognitive Examination (Hsieh et al., 2013; Mathuranath et al., 2000). In the original VOSP, these letters are presented at a fixed level of degradation (around 30% “completeness”), leading to ceiling effects in typical adults and early-stage PCA patients (Firth et al., 2019), and limiting the sensitivity of the test in tracking disease progression. The Graded Incomplete Letters Test (GILT; Yong et al., 2024) was developed to fill this gap by measuring the threshold for recognising letters under increasing degradation (decreasing their “completeness”, as shown in Figure 1). Typical adults (such as those in the UK Biobank) require completeness levels around 5-6% to reach threshold on this test, while PCA patients demonstrate significantly poorer performance, with a median completeness threshold around 47% (Yong et al., 2024). Although the GILT can therefore clearly differentiate PCA patients from cognitively typical adults, the underlying mechanisms are poorly understood. In particular, it is unclear which of the cortical symptoms that arise in PCA could drive these deficits in the recognition of incomplete letters.

**Figure 1.**
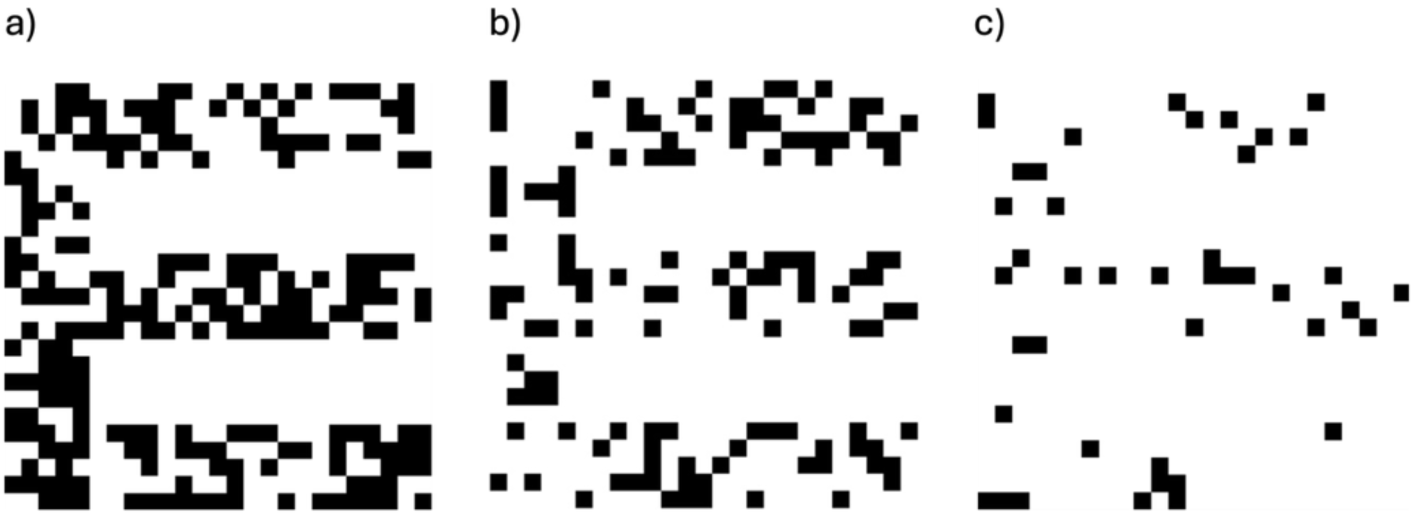
Example stimuli from the Graded Incomplete Letters Test. Each panel shows the letter “E” at different “completeness” levels, defined as the amount of black “ink” within the letter that is present relative to the white background. Completeness levels are a) 0.5 proportion complete, close to typical thresholds in people with PCA, b) 0.3, close to stimuli used in the VOSP battery and b) 0.1, closer to typical thresholds.

A number of the symptoms exhibited by PCA patients could drive impairments in GILT performance. A strong candidate mechanism is crowding, the impaired recognition of objects within cluttered environments (Levi, 2008). Crowding is a low-to-mid level cortical process in typical vision (Anderson et al., 2012; Flom et al., 1963; Levi et al., 2002) that has a significant impact on object and letter recognition in visually cluttered environments (Levi, 2008; Levi et al., 2002; Pelli et al., 2004). In individuals with typical vision, crowding predominantly occurs in peripheral vision (Bouma, 1970; Pelli et al., 2004), with far smaller effects in foveal/central vision (Coates et al., 2018). In PCA, crowding has been shown to be elevated in the fovea (Crutch & Warrington, 2007; Crutch et al., 2016; Strappini et al., 2017; Yong, Shakespeare, Cash, Henley, Nicholas, et al., 2014). In the typical periphery, these disruptions occur independently of visual acuity – an object may be sufficiently large to be recognized in isolation, but difficult to identify when surrounded by other stimuli (Levi, 2011; Pelli et al., 2004). Though crowding is typically observed when the recognition of a target object/letter is impaired by surrounding flanker object/letters (Levi et al., 2002; Pelli et al., 2004), crowding can also occur internally within a single object. Internal crowding has been observed in face and word recognition (Martelli et al., 2005) as well as within single Chinese characters, where their complex internal structure leads to “within-character” crowding (Zhang et al., 2009). In the context of the GILT, although only one letter is presented, their composition of spatially separated fragments may give rise to internal crowding between the fragments/features, potentially contributing to the recognition deficits observed in patients with PCA.

Another candidate symptom likely to impair GILT performance is global integration, a high-level cortical process where individual visual features are combined into a coherent object (Loffler, 2008; Prinzmetal, 1981). Here, we focus on the spatial integration of letter components (such as edges, corners/junctions, and strokes), as opposed to broader conceptions of feature integration involving the binding of visual attributes (like colour and motion; Treisman, 1988). In typical vision, incomplete letter recognition has been shown to depend upon both the availability of diagnostic letter components (e.g. junctions and line terminations) and their integration across space and time (Fiset et al., 2008; Grainger et al., 2008; Greene & Visani, 2015; Pelli et al., 2009). These integrative processes become particularly important when local features are degraded through brief presentations or local feature masking (Di Lollo et al., 1994; Hogben & di Lollo, 1974; Zhang et al., 2024). Impairments in global integration have been argued to be a core feature of PCA, based on the early onset of difficulties with recognising letters, words and scenes, particularly when degraded or shown with non-canonical presentation such as odd angles or cursive letter fonts (Coslett & Saffran, 1991; Mendez et al., 2007; Yong, Shakespeare, Cash, Henley, Warren, et al., 2014). Integrative deficits may similarly underlie reported difficulties in contour integration (Uhlhaas et al., 2008), visual search and distractor suppression tasks (Mendez & Cherrier, 1998), as well as difficulties with face recognition, which requires the holistic integration of facial features (Crutch et al., 2012), and issues reproducing the Rey-Osterrieth Complex Figure (Li et al., 2018). The documented patterns of atrophy in occipital regions in PCA patients may drive these global integration deficits, given the importance of extrastriate areas for global-form perception (Loffler, 2008). In the context of the GILT, where letters are degraded with spatially separated fragments, successful recognition depends crucially on the integration of these fragmented features (e.g. edges, corners, and strokes) into a coherent letter, which these integration deficits would impair.

Although we assume that incomplete letter recognition relies predominantly on cortical processes, it is nonetheless important to rule out limitations from optical factors such as blur or reduced contrast sensitivity. Since visual acuity is often preserved in the early stages of PCA (Benson et al., 1988; Crutch et al., 2017), it is unlikely that PCA patients would experience basic visibility challenges with the large stimuli used in this test. However, given that PCA patients are typically older adults (with an average onset age of 59 years old; Chapleau et al., 2024; Crutch et al., 2012; Schott et al., 2016), there is an increased likelihood of age-related ocular conditions such as cataract or glaucoma, which can reduce visibility through increased blur and decreased contrast sensitivity (Christie et al., 2012; Stamper, 1984). These optical factors could contribute to poorer performance on the GILT, potentially making it more difficult to isolate cortical deficits within this age group (Harding et al., 2018; Yong et al., 2024) or leading to misdiagnoses. Establishing the relative contributions of cortical and optical factors is therefore essential for interpreting deficits in GILT performance.

To better understand the mechanisms underlying GILT performance, we sought to simulate the clinical visual deficits of people with PCA in healthy adults. This approach has been widely used in previous research on other conditions, for example with amblyopia simulated using peripheral vision (Kalpadakis-Smith et al., 2022) and nystagmus simulated through stimulus motion (Tailor et al., 2021). Simulation of this kind allows these factors to be systematically isolated and manipulated in ways that would be infeasible with clinical populations, due to the length of testing required and/or task demands. Here, we employed this simulation approach to test the limits of performance on the GILT, focusing on the effect of the above candidate optical and cortical factors. The investigation was divided into three main experiments. In Experiment 1, we examined whether poor performance on the test could be attributed to optical issues by introducing blur and low contrast. In Experiment 2, we examined whether the deficits in GILT performance were associated with cortical issues by simulating the effects of elevated crowding and difficulties in global integration across space. Finally, in Experiment 3, we further explored cortical limitations by simulated elevated crowding and difficulties in temporal global integration. Based on the above evidence regarding the nature of the deficits in PCA, we hypothesized that impaired GILT performance would be driven primarily by cortical factors, such as elevated crowding and impaired global integration, whereas optical factors such as blur and low contrast would have negligible effects.

### Experiment 1: Optical factors

Our first aim was to determine whether optical issues impair performance on the GILT. Although optical factors affect the visibility of letters in general (Westheimer, 2016), their influence on thresholds for incomplete letter recognition is unknown. We approached this by simulating the effects of these optical factors in typical adults by manipulating the level of blur and contrast applied to incomplete letter stimuli.

#### Method

##### Design

To rule out the possibility that optical factors could influence GILT performance through basic visibility limitations (i.e. letter detection), we first measured detection thresholds for blur and contrast using complete letters. We then examined how these optical factors influence performance on the GILT by measuring thresholds for letter completeness using stimuli at five levels of blur or contrast, all of which exceeded the respective detection thresholds.

##### Participants

Six participants (age range: 20–30 years, 5 females, 1 left-handed, 1 left dominant eye) with normal or corrected-to-normal visual acuity participated in the study, including author ZH. All participants reported that they did not have diagnoses of dyslexia or any developmental or age-related visual conditions like amblyopia or glaucoma. The dominant eye was tested using the Crider ring test (Crider, 1944). All participants gave written informed consent prior to participation. Procedures were conducted in accordance with the principles of the Declaration of Helsinki and approved by the UCL Experimental Psychology Research Ethics Committee.

##### Apparatus

Stimuli were presented on a 32-inch wide-screen Display++ monitor (Cambridge Research Systems Ltd.) at a resolution of 2560×1440 pixels and 120 Hz refresh rate. The monitor was calibrated with a Minolta LS110 photometer to give a minimum luminance of 0.16 cd/m^2^ and a maximum of 143 cd/m^2^. Participants were seated in front of the monitor in a dark testing room, with stimuli viewed binocularly at a distance of 60 cm from the screen. Head movements were minimised using forehead and chin rests. A PowerMate response dial (Version 3.1.0; Griffin Technology) was used to record responses.

##### Stimuli

Stimuli were generated using MATLAB (MathWorks; version R2020b) and presented using an Apple iMac running Psychtoolbox (version 3.0.18; Brainard, 1997; Pelli, 1997). Stimuli were one of 12 uppercase letters from the set “C D E F H K N P R U V Z”, presented in an extended Sloan font (see Figure 2; Sloan, 1959), as in the standard GILT protocol (Yong et al., 2024). Sloan letters are a standard in acuity testing because of their fixed letter proportions, in which all structural features are scaled relative to letter size such that the stroke width corresponds to one-fifth of the letter diameter. This format ensures that during the experiment the effects of degradation on the letter are equivalent across different letterforms. The pixelated “checks” used in incomplete letters were similarly scaled to be one-fifth of each stroke width, meaning that each stroke was composed of five checks that could be removed as degradation increased. Letters were shown at a width of 8.6 degrees of visual angle, which is similar to the stimulus size used by Yong et al. (2024). All stimuli were presented on a uniform grey background.

**Figure 2.**
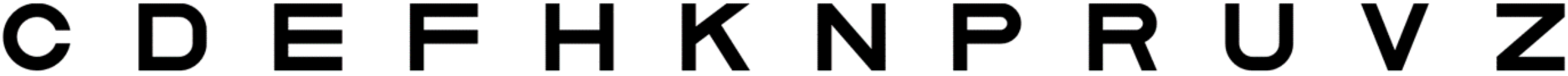
The 12 potential letterforms from the extended Sloan font used as incomplete letter stimuli, here shown without degradation.

To exclude the possibility that our manipulations of blur and low contrast impaired performance due to simple visibility issues (which would affect the recognition of complete/undegraded letters as well), initial measurements of blur and contrast thresholds were conducted using complete letters with adaptively-applied levels of blur or contrast. Sample stimuli are shown in Figure 3.

**Figure 3.**
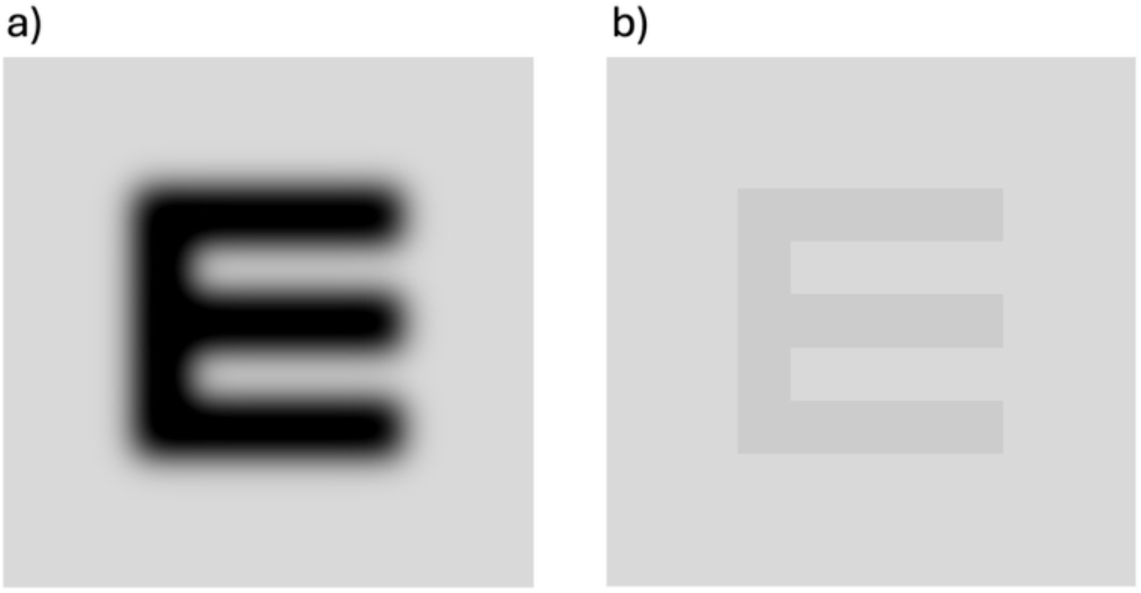
Complete letter stimuli with blur or lowered contrast applied. Examples of the letter “E” with full completeness under conditions of a) blur (cut-off at 5 cycles/image) and b) low contrast (10%).

The main experiment had 3 stimulus types. In addition to the standard GILT stimuli, two additional manipulations were tested separately: blurred stimuli and reduced-contrast stimuli. Blurred stimuli were generated by transforming the images into the Fourier domain and applying a low-pass filter with cut-off frequencies of either 1.75, 2.5, 5, or 10 cycles/image (0.2, 0.3, 0.6 or 1.2 cycles/deg). The chosen range was selected based on pilot testing to produce measurable effects while remaining above visibility thresholds and covering the spatial-frequency range most relevant for letter recognition (approximately 3 cycles/letter; Solomon & Pelli, 1994). For contrast manipulation, letters were presented at Weber contrast levels of 3, 5, 9, and 18%, likewise chosen to span a range of perceptual effects whilst still remaining above visibility thresholds. Sample incomplete-letter stimuli are shown in Figure 4.

**Figure 4.**
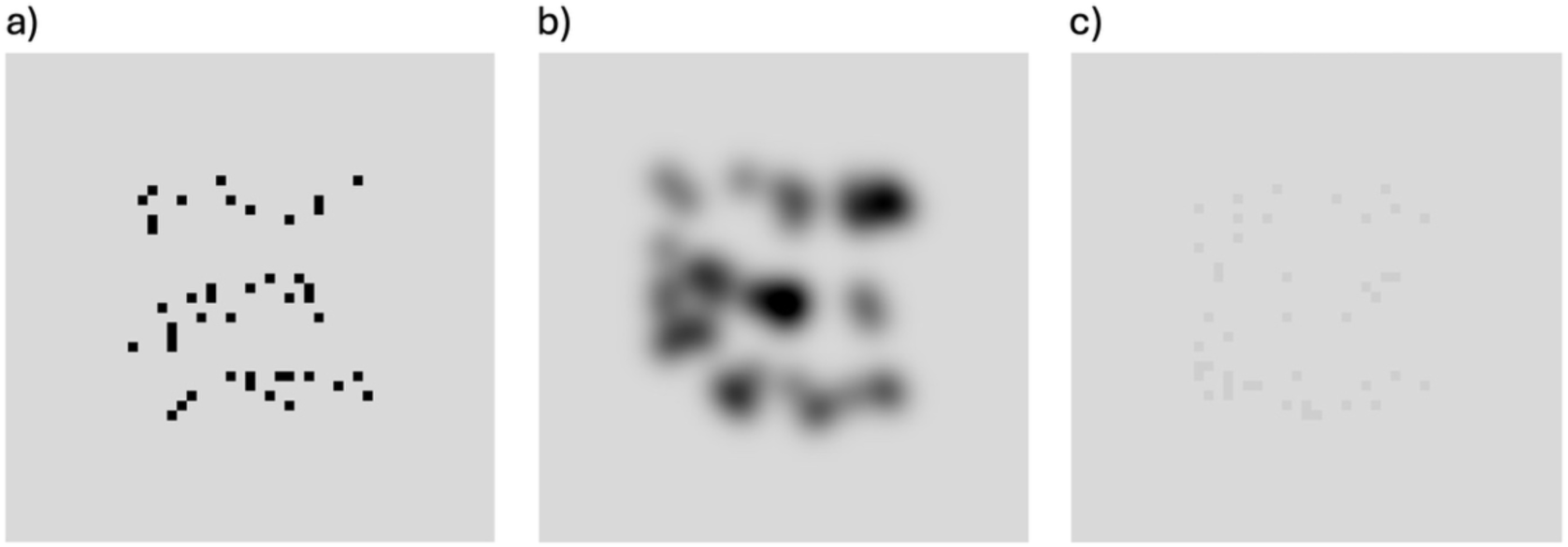
Incomplete letter stimuli with blur or lowered contrast applied. Examples of the letter “E” with the same completeness level (0.1 proportion complete), either a) at baseline, as in the standard GILT, b) with blur (5 cycles/image), or c) with low contrast (9%).

##### Procedure

To ensure that stimuli in the main experiment were perceptually detectable, we first measured individual detection thresholds for blur and contrast using complete letters. These measurements were conducted prior to the main task. Thresholds were estimated using an adaptive QUEST procedure (Watson & Pelli, 1983). Each trial involved presenting a fully complete letter with a given blur or contrast level, which over trials was varied adaptively to converge on the threshold corresponding to 54.2% accuracy (midway between chance and ceiling). To avoid rapid convergence of the QUEST, we added variance to the blur/contrast levels presented on each trial by adding a value selected from a Gaussian distribution (with 0 mean and 1 SD) multiplied by 0.3× the current threshold estimate. This minimised the number of trials presented at the same level, which improves the subsequent fit of psychometric functions to the data, as in prior work (Tailor et al., 2021). There were 55 trials for each block, including 5 practice trials at the beginning of each block that were not used for analysis. The remaining 50 trials included 5 catch trials to ensure attention was maintained throughout (and to facilitate psychometric curve fitting), where letters were presented with no blur and full contrast for the 5^th^, 15^th^, 25^th^, 35^th^, and 45^th^ trials. Each block was repeated three times. These measurements were used to ensure that the fixed blur and contrast levels employed in the main experiment were above each participant’s detection thresholds, such that all stimuli used to measure completeness thresholds remained detectable.

In the main experiment, completeness thresholds for 5 different levels of blur and contrast were measured (four levels of manipulation plus a baseline). Stimuli were degraded to varying levels of completeness, which was adjusted based on participants’ performance to converge on the threshold level (54.2% correct) using the same adaptive QUEST procedure as described above, though here applied to completeness levels rather than blur/contrast. This included the added variance (again at 0.3× the threshold estimate on each trial) and catch trials (presented at full completeness). Each block of trials was again 55 trials (including 5 initial practice trials presented at 0.3 proportion complete), with each block repeated three times, yielding 30 blocks in total (15 blur and 15 contrast). To reduce confusion for participants, the blur and contrast manipulations were conducted separately during the experiment, each with its own baseline condition with no blur and full contrast (similar to the standard GILT). The order of conditions was randomised between participants, with the blur experiment conducted first for three participants and contrast first for the remainder. Prior to the start of the experiment, participants completed brief practice blocks to familiarize themselves with the task. During these practice blocks, five different levels of blur or contrast were presented. The whole experiment took approximately 3.5 hours, split over 2-3 sessions.

For both the detection threshold experiment and the main completeness threshold experiment, each trial began with participants viewing a white Gaussian fixation point at the centre of the screen. A target upper-case letter stimulus was then displayed foveally for 500 ms, immediately followed by a 500 ms visual mask, to prevent continued cortical processing of the stimulus after offset (Breitmeyer & Ogmen, 2006). The mask consisted of a random-check noise pattern with the same dimensions as the letter stimulus, including the spatial structure (i.e., noise checks matched the size of checks within the letters), with pixel intensities drawn from a Gaussian distribution of grey levels. Subsequently, twelve lower-case letter options appeared in a ring at the centre of the screen (a 12-alternative forced choice response method; 12-AFC), with participants asked to rotate the response dial to indicate the target letter they saw, selecting the corresponding lower-case letter from the 12-letter set. As the dial was turned, the ring of response options rotated accordingly, with the current response shown as the largest letter. Participants were instructed to respond as accurately as possible, with response letters remaining visible until the response was made. When uncertain about the letter, they were required to guess. After each response, a 500 ms inter-trial interval was maintained before presenting the subsequent uppercase letter. Throughout the run, participants were instructed to keep fixating on the fixation dot. An example trial is shown in Figure 5.

**Figure 5.**
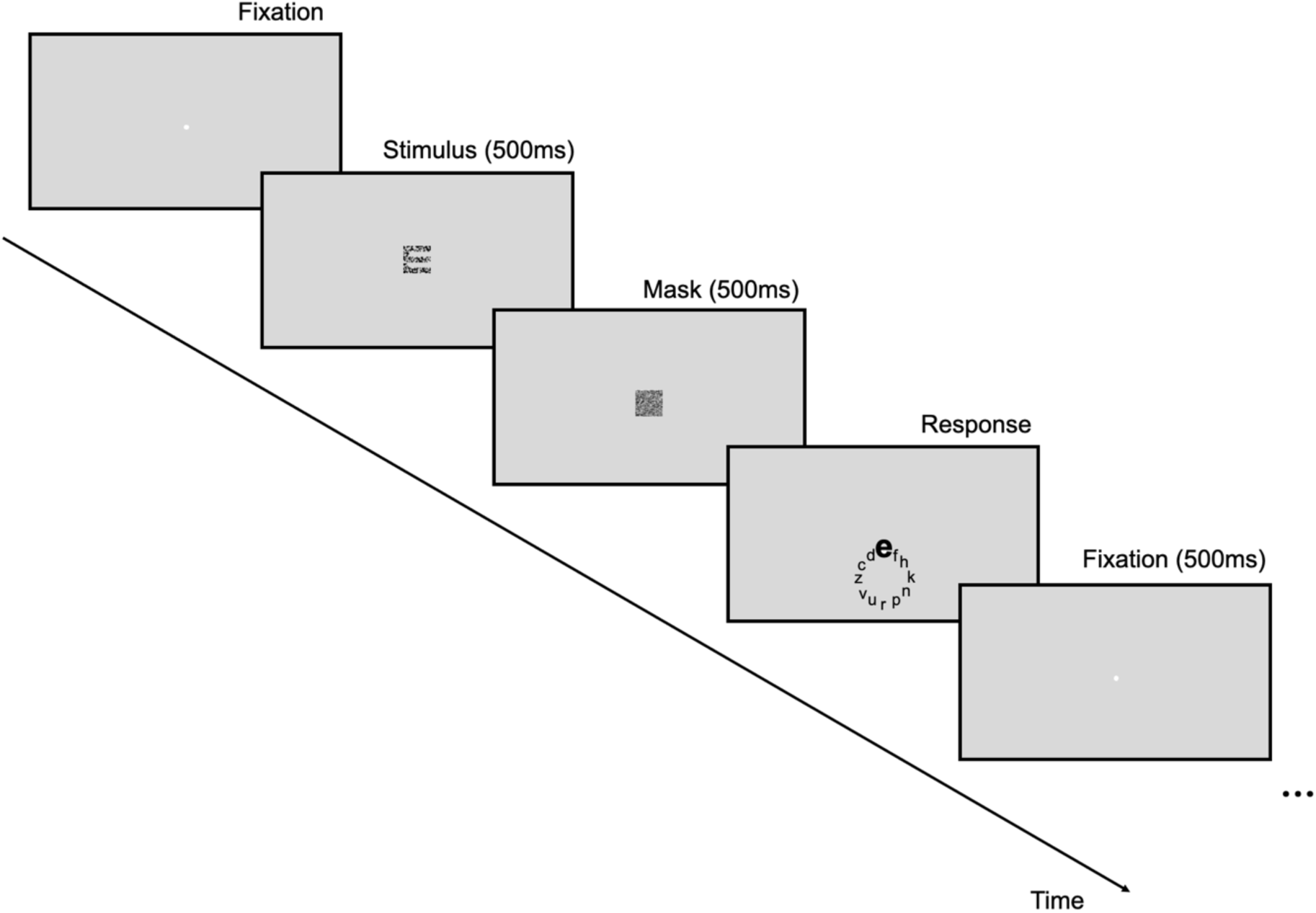
An example trial in Experiment 1 for the main completeness thresholds experiment. After an initial fixation period, an incomplete letter stimulus was presented at a given level of completeness (determined by the QUEST procedure), followed by a mask and the response period. A standard GILT stimulus is depicted for this trial, as in the baseline conditions.

##### Data analysis

Data were collated and analysed in MATLAB. For detection thresholds, blur and contrast sensitivity were analysed separately, with responses pooled across trials for each participant to estimate threshold. Blur thresholds were defined in cycles/image and contrast thresholds as Weber contrast. For GILT thresholds (expressed as the proportion complete), responses from repeated blocks were again combined, separately for each participant and stimulus condition (baseline, blur and contrast), to give 150 trials per stimulus condition. Completeness level values were binned together with a resolution of 0.01 proportion complete, with the proportion of correct responses calculated for each bin. Cumulative Gaussian functions were then fitted to this data using two free parameters (mean and variance) using weights adjusted by the number of trials at each completeness level. As above, thresholds for each condition were taken as the level of blur, contrast, or completeness at which accuracy reached 54.2%. Figure 6 presents example data for one participant across 3 blur conditions of the main completeness threshold experiment. To analyse the main experiment, two-way mixed-effects analyses of variance (ANOVAs) were conducted separately for the blur and contrast conditions, with fixed effects of blur or contrast, and a random effect of participant (Howell, 1997, p.485). This mixed-effects approach was used to account for the repeated-measures structure of the design and to explicitly model between-participant variability.

**Figure 6.**
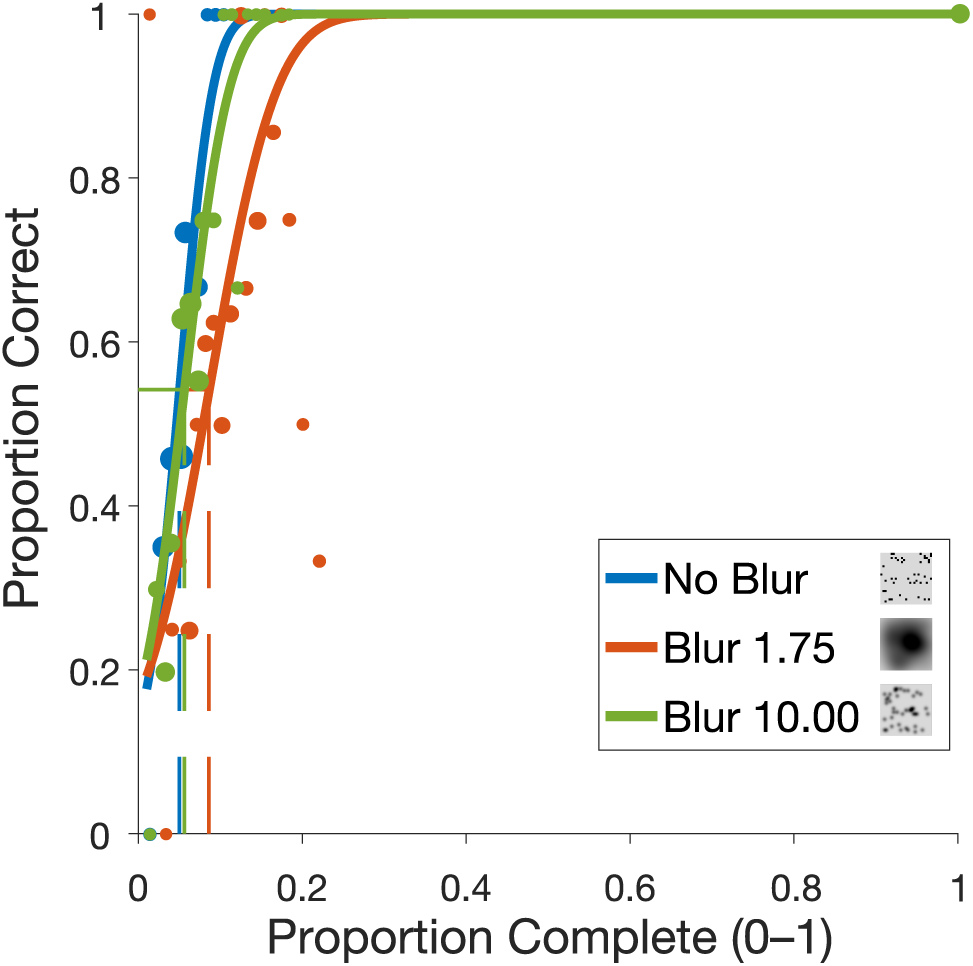
Example data from a single participant in the main completeness experiment. The proportion of correct responses is shown across varying levels of letter completeness (expressed as proportion complete) for three example blur conditions: no blur (blue), 1.75 cycles/image (orange), and 10 cycles/image (green). Dashed coloured lines represent completeness thresholds, taken at the completeness level where performance reached 54.2%.

#### Results

Completeness thresholds (the proportion of letter completeness required to reach 54.2% correct) are shown in Figure 7 at different levels of blur and contrast. Figure 7a shows the effects of blur. Blur thresholds for the detection of complete letters are shown by the red dotted line, showing that the average blur threshold was 1.148 cycles/image. This threshold reflects the spatial frequency cut-off at which a complete letter could be detected reliably, which served as a minimum for selecting blur levels in the experiment – all were above this detection threshold to ensure visibility of the incomplete letters.

**Figure 7.**
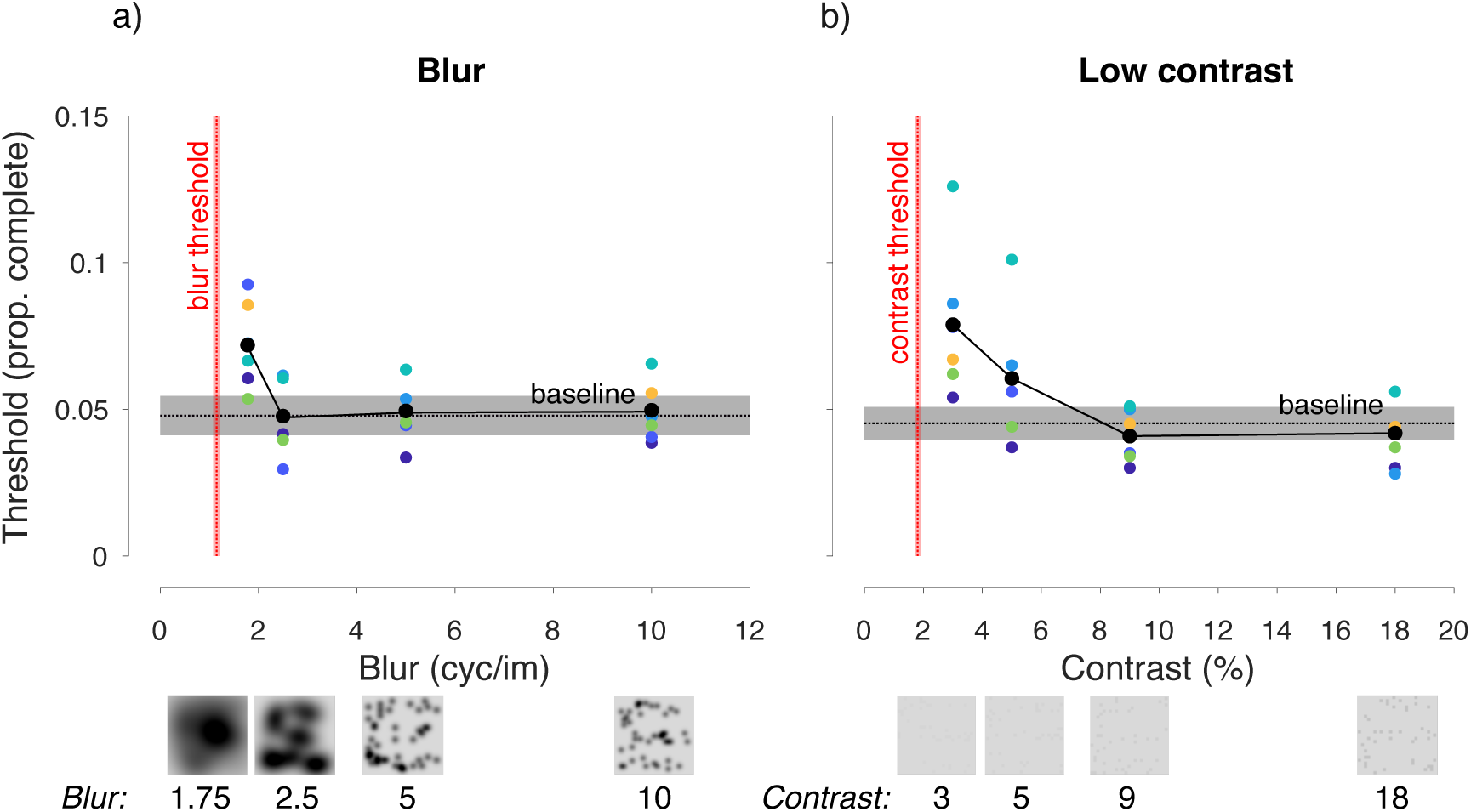
The effect of optical factors on completeness thresholds (Experiment 1). a) Effects of blur. Black dots show the mean completeness threshold (expressed as proportion complete) for each blur condition, plotted as a function of the filter cut-off applied to stimuli (in cycles/image); coloured dots plot individuals. The black horizontal dashed line plots the mean threshold of the baseline condition, with the grey area showing the SEM. The red vertical dashed line shows the average blur threshold for complete letters, with the red shaded area showing the SEM. Example stimuli at each blur level are shown along the x-axis with 0.1 completeness. b) Effects of contrast, with completeness thresholds plotted as a function of Weber contrast, following the same conventions as a), except that the vertical red solid line shows the average contrast-detection threshold for complete letters.

For completeness thresholds, baseline values without blur applied averaged around 0.05 proportion complete (i.e. 5% of the complete letter), consistent with the results for typical adults in Yong et al. (2024). With blur applied, thresholds were indistinguishable from the baseline across most of the blur range. It is only at the lowest cut-off value of 1.75 cycles/image (with the highest blur) that small threshold elevations up to 0.08 proportion complete were evident, indicating that participants needed slightly higher ‘completeness’ (i.e., more ‘ink’ or visible pixels) to recognise the letters under this condition. A two-way mixed-effects ANOVA confirmed this pattern, with a significant main effect of blur, *F*(4,20) = 7.13, *p* = 0.001. The analysis also revealed a significant main effect of participant, *F*(5,20) = 4.7, *p* = 0.005, indicating inter-individual variability in performance. Nonetheless, contrasts show that thresholds at 1.75 cycles/image were significantly elevated relative to baseline, *t*(5) = –3.087, *p* = 0.027. At higher cut-off frequencies (2.5, 5, and 10 cycles/image), thresholds did not differ significantly from baseline (all *p* > 0.05). Thus, performance was only mildly affected by blur, and only at levels very close to the visibility limit for complete letters.

Figure 7b shows the effect of contrast on completeness thresholds. Detection thresholds for complete letters averaged 1.8% Weber contrast, shown by the red dotted line, with all contrast conditions in the main experiment shown above this detection level. At full contrast, the baseline completeness threshold was approximately 0.05, similar to the blur experiment. Thresholds were again close to baseline for much of the contrast range, rising only slightly when contrast was reduced near to the detection limit, where thresholds reached a peak around 0.08 completeness. This slight decline in recognition performance was confirmed by a two-way ANOVA, with a significant main effect of contrast, *F*(4,20) = 4.01*, p* < 0.001. The main effect of participant was also significant, *F*(5,20) = 10.02, *p* < 0.001, indicating between-participant heterogeneity. Contrasts show that thresholds were nonetheless significantly elevated at 3% *(t*(5) = –5.439, *p* = 0.003) and 5% *(t*(5) = –3.271, *p* = 0.022) relative to baseline, whereas thresholds at 9% and 18% did not differ significantly (both *p* > 0.05). As shown in Figure 7b, recognition performance returned to baseline as contrast increased and remained stable across most of the contrast range, indicating that contrast only mildly impaired performance near the visibility limit.

Overall, the results of Experiment 1 indicate that incomplete letter recognition was only slightly impaired at extremely high levels of blur and low levels of contrast that fell near to the detection thresholds for complete letters. Even in these instances, completeness thresholds were only mildly elevated to reach levels around 0.08 completeness. Importantly, even at these reduced performance levels, recognition remained substantially better than that observed in patients with PCA, whose median threshold was 0.47 (Yong et al., 2024). Moreover, even small reductions in blur or increases in contrast caused the effect of these optical factors to drop away completely, suggesting performance on the GILT is largely unaffected by blur or contrast reduction, even at moderate-to-high levels of each. These findings therefore support the hypothesis that optical factors alone cannot account for the levels of poor incomplete letter recognition observed in PCA. Consequently, we next investigated whether incomplete letter recognition is more strongly associated with cortical factors.

### Experiment 2: Crowding and spatial integration

To investigate whether impaired performance on the GILT is linked to cortical processing difficulties, we focussed on two processes likely to limit incomplete letter recognition that are known to be abnormal in PCA – crowding (Crutch & Warrington, 2007; Crutch et al., 2016) and global integration (Metzler-Baddeley et al., 2010). Crowding was manipulated by presenting stimuli at increasing eccentricities in the peripheral visual field, where crowding effects are increased in typical vision (Bouma, 1970; Pelli et al., 2004). Global integration demands were modulated by varying check sizes, which affect how spatial information is distributed across each letter – smaller checks spread the available features more evenly, increasing the likelihood that critical elements remain visible, while larger checks cause the “ink” to concentrate in fewer areas, leading to the more rapid loss of key features. With these manipulations, we can thus vary both the visibility of individual features and how effectively participants could integrate local visual features into coherent letter representations.

#### Method

##### Design

To investigate the role of these two cortical factors in incomplete letter recognition, we measured thresholds for letter completeness on the GILT using stimuli with three visual field locations and four check sizes.

##### Participants

Six participants took part (age range: 20-24 years, 5 females, all right-handed, 1 left dominant eye), including author ZH and two participants who participated in Experiment 1, with the rest newly recruited. The participant criteria, ethical considerations, and approval process for Experiment 2 were identical to those described for Experiment 1.

##### Apparatus

The set up for Experiment 2 was largely similar to that described for Experiment 1. In addition to the apparatus used in Experiment 1, an EyeLink 1000 (SR Research, Mississauga, ON, Canada) was used to monitor participants’ fixation during trials with stimuli in peripheral vision.

##### Stimuli

Stimuli were generated as in Experiment 1. To examine the role of global integration in incomplete letter recognition, we varied the width of the checks within each stroke. Stimuli were shown with 2, 3, 5, and 10 checks per stroke, where 5 checks/stroke is the baseline level used in Experiment 1 as well as the standard GILT (Yong et al., 2024). These manipulations alter the distribution of the information removed from local features such as strokes and junctions, thus varying the ability to integrate these features into coherent global forms (Grainger et al., 2008; Greene & Visani, 2015). Sample stimuli are shown in Figure 8. Stimuli were displayed at a size of 8.6°, except for the 3 checks per stroke condition, which was slightly smaller (8.41°) due to the need to render stimuli using integer pixel values; pilot testing confirmed that this difference had no measurable impact on performance. The stimuli were presented at one of three locations: the fovea, and peripherally at 10° and 20° eccentricity in the upper visual field. Given that crowding primarily occurs in peripheral vision (Bouma, 1970; Pelli et al., 2004), by presenting stimuli peripherally in healthy participants, we aimed to simulate the elevated foveal crowding observed in PCA patients. In addition, because PCA is associated with both impaired global integration and elevated crowding, we also sought to examine the combined effects of these impairments on incomplete letter recognition. The overall design therefore included 3 (eccentricity: fovea, 10°, 20°) × 4 (check size: 2, 3, 5, 10 per stroke) conditions, enabling us to assess the effects of crowding and global integration on letter recognition performance both independently and in combination.

**Figure 8.**
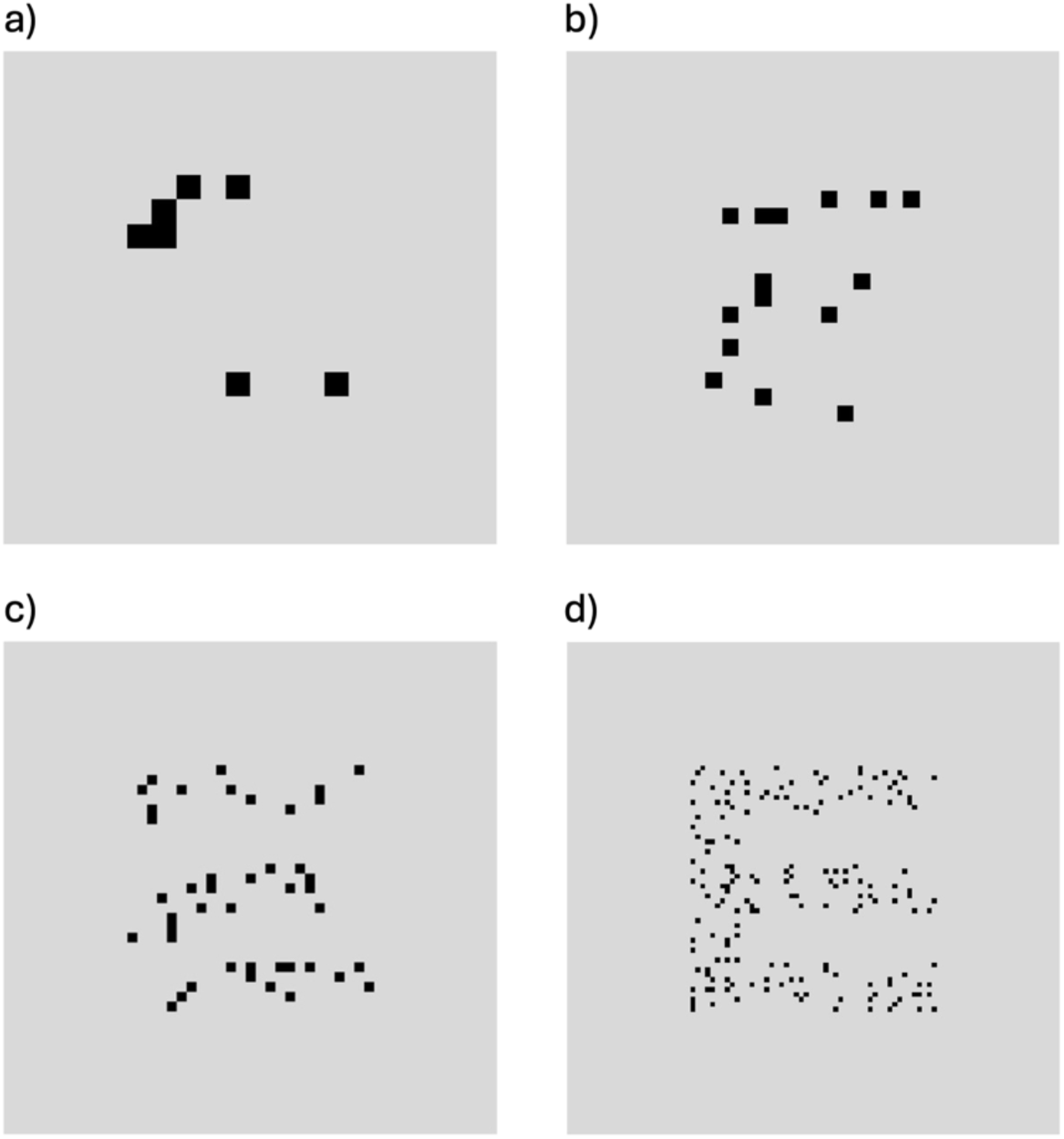
Example incomplete letter stimuli with different check sizes. The letter “E” is shown with different check sizes (quantified as checks per stroke) at the same completeness level of 0.1. a) 2 checks/stroke b) 3 checks/stroke c) 5 checks/stroke (the standard GILT baseline) d) 10 checks/stroke.

##### Procedure

To begin the experiment, the eye tracking equipment was set up and calibrated using the built-in 5-point calibration. In both foveal and peripheral conditions, participants were instructed to fixate on the fixation dot (near the bottom of the screen) during stimulus presentation. Fixation was not required during the response phase. For each trial to begin, fixation had to be within a 1.5° radius around the fixation point. If fixation moved outside this window, the trial was immediately cancelled, and both a sound alert and a message appeared on the screen to remind participants to fixate on the dot. Once the participant fixated, a new trial would begin. Re-calibration of the EyeLink was performed if fixation was lost for over 5 sec or on more than 20 trials, and at least once every 15 minutes.

The experiment consisted of twelve conditions (4 check sizes × 3 eccentricities). Each block of trials was run using the same QUEST procedure as Experiment 1, which varied completeness levels to find threshold at 54.2% correct. Stimulus location and check size was held constant within each block of trials. There were 55 trials for each block, including 5 practice trials at the beginning of the task (not used in analyses) and 5 catch trials across each block. Each condition was repeated three times to give 36 blocks in total. The whole experiment took approximately 3.5 hours, split over three sessions.

##### Data analysis

Data were analysed in MATLAB. Completeness thresholds were determined for each condition following the same procedure as in Experiment 1. A three-way mixed-effects analysis of variance (ANOVA) was conducted, with fixed effects of check size (2, 3, 5, and 10 checks per stroke), eccentricity (0°, 10°, and 20°), their two-way interaction, and a random effect of participant (Howell, 1997, p.485).

#### Results

Completeness thresholds, expressed as the proportion complete required to reach 54.2% accuracy, were computed for different visual-field locations and check size, as shown in Figure 9. As in Experiment 1, smaller completeness values indicate that participants were able to identify letters with fewer visual cues. Thresholds systematically increased with eccentricity, with poorer performance in peripheral vision compared to the fovea, with a significant main effect of eccentricity: *F*(2,71) = 61.58, *p* < 0.001. Varying check size also significantly influenced performance, with a significant main effect of check size: *F*(3,71) = 73.51, *p* < 0.001. Performance was best (i.e. thresholds were lowest) with the finest-scale checks (10 checks/stroke), and decreased as checks became larger, with the worst performance at 2 checks/stroke. Importantly, there was a significant interaction between eccentricity and check size, *F*(6,71) = 14.92, *p* < 0.001, such that performance with the larger checks (e.g. 2 checks/stroke) was disproportionately impaired as eccentricity increased. In the fovea, average thresholds with 5 checks/stroke (the standard GILT baseline) were 0.036, slightly lower than those in Experiment 1 and Yong et al. (2024). Relative to this point, the lowest threshold was 0.007 completeness with 10 checks per stroke in the fovea, rising to the highest threshold level of 0.23 with 2 checks/stroke at 20° eccentricity. In addition, there was a significant main effect of participant, indicating clear between-participant heterogeneity, *F*(5,71) = 4.94, *p* = 0.007. The interaction between check size and participant was also significant, *F*(15,71) = 3.19, *p* = 0.003, suggesting individual variability in sensitivity to check size. There was no significant eccentricity × participant interaction (*F*(10,71) = 1.69, *p* = 0.131), indicating that the influence of eccentricity on performance was consistent across individuals.

**Figure 9.**
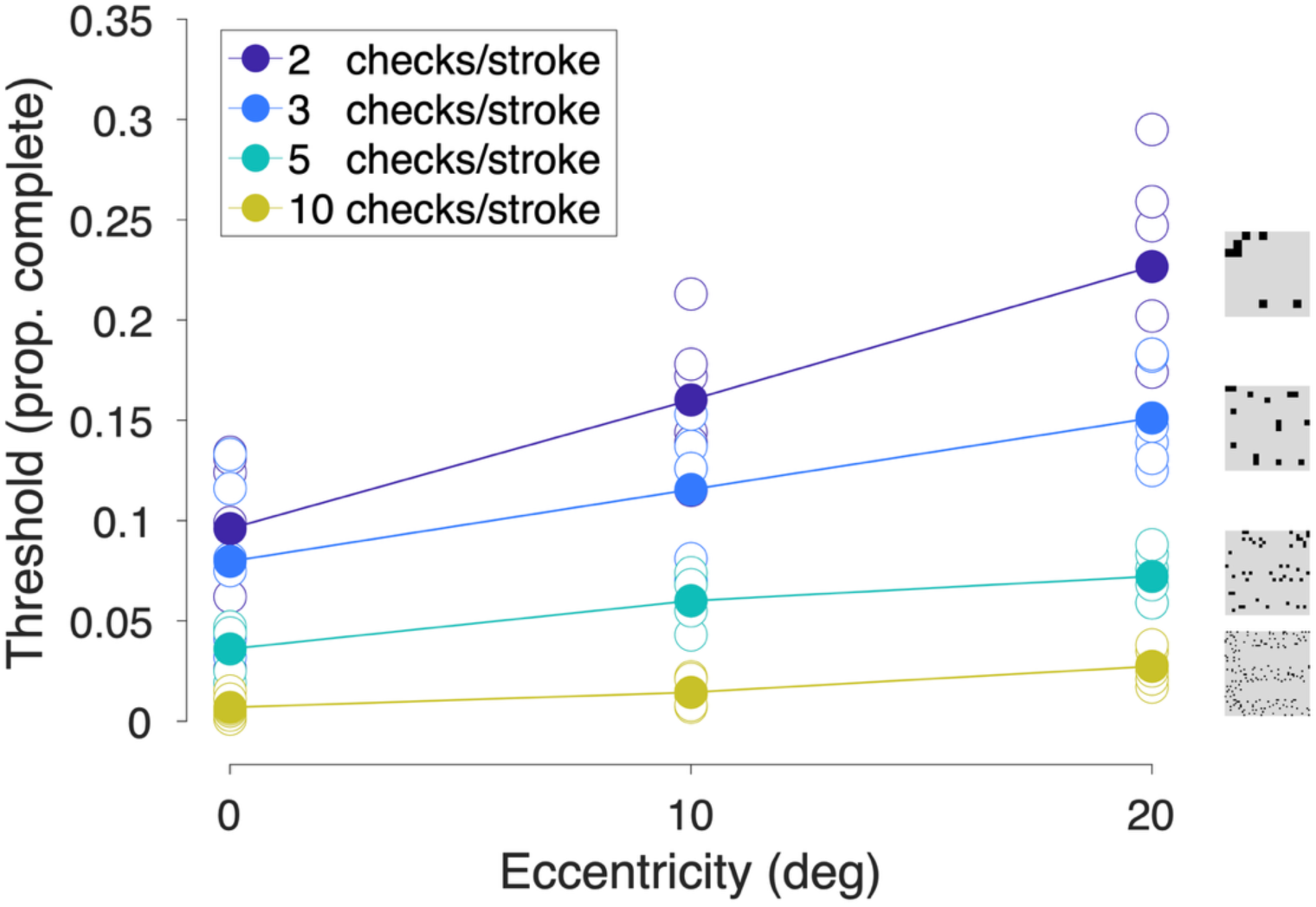
Completeness thresholds from Experiment 2 plotted as a function of eccentricity, separately for stimuli with different check sizes (see legend). Solid dots show the mean threshold for each condition; hollow dots represent the threshold for each participant. Example stimuli with each check size are shown on the right-hand side.

Overall, the findings of Experiment 2 suggest that poor incomplete letter recognition is more likely associated with cortical than optical factors, as variations in both the level of crowding and the ease of global integration strongly affected performance, particularly through their interaction. The most challenging condition was where stimuli with 2 checks/stroke were shown at 20° eccentricity, which combines the high crowding of peripheral vision with concentrated degradation of the letters (i.e. each drop in completeness involved the removal of ‘ink’ from a smaller number of larger regions, relative to other conditions where the degradation was more distributed). This combination gave average completeness thresholds of 0.23, which more closely approximates the levels observed in PCA patients (median: 0.47; Yong et al., 2024), compared to the results of Experiment 1. In the following experiment, we sought to further test the hypothesis that performance is limited by cortical factors with another manipulation intended to impair global integration.

### Experiment 3: Crowding and temporal integration

While the previous experiment manipulated global integration by altering the spatial properties of letters, in Experiment 3 we used temporal variations of the features within the incomplete letter stimuli, along with limited lifetime presentation. This temporal manipulation was used to engage mechanisms of sequential global integration, requiring the visual system to combine information across time (Di Lollo et al., 1994; Hogben & di Lollo, 1974), which has previously been shown to strongly impair letter recognition (Greene & Visani, 2015; Zhang et al., 2024). This approach therefore provides an alternative method for assessing global integration processes within the GILT framework. By presenting stimuli in the peripheral visual field, where crowding effects are increased, the experiment also allowed us to examine how this temporal manipulation interacts with crowding to influence recognition performance. Although temporal differences between target and flanker elements can reduce crowding in some contexts (Chakravarthi & Cavanagh, 2007; Greenwood et al., 2014; Huckauf & Heller, 2004), this was unlikely in our study because the letter fragments appeared and disappeared rapidly in an interleaved sequence, rather than being systematically segregated. By interspersing the transient onset signals associated with the letter features, these temporal variations should not therefore reliably separate the constituent features or reduce within-letter crowding (Greenwood et al., 2014). Instead, this dynamic manipulation allowed us to examine the conjunction of both temporal integration limits and crowding effects.

#### Method

##### Design

As in Experiments 1 and 2, letter completeness thresholds were measured using the GILT, here with stimuli presented at three visual field locations and under two presentation modes: static or dynamic.

##### Participants

Six participants took part (age range: 22-24 years, 5 females, all right-handed, all right dominant eye), including authors ZH and two participants who participated in Experiments 1 and 2, with the rest newly recruited. Participant criteria and ethical approvals were identical to those of Experiment 1.

##### Stimuli and Procedure

The apparatus for Experiment 3 was identical to that of Experiment 2. Stimuli were largely similar to those of Experiment 2, with the key difference that the stimuli were shown either in static (i.e., baseline) or in dynamic (temporally extended) form. This manipulation aimed to investigate the role of temporal integration in incomplete letter recognition by varying the presentation of visual information over time.

Both static and dynamic conditions used the standard GILT stimuli with a check size of 5 checks/stroke. In the dynamic condition, each stimulus consisted of 25 frames presented sequentially at 25 ms per frame, giving a total duration of 625 ms. Each individual check had a lifetime of 75 ms and was therefore visible across three consecutive frames. These parameters were determined through pilot testing to establish values that maintained task difficulty (by distributing features across a sufficiently long temporal period), with a refresh rate below the flicker fusion threshold (Skrandies, 1985) to ensure sequential frames were not perceptually fused, whilst also ensuring the visibility of individual checks. On each frame, 25 of the 625 possible checks were newly revealed, which could either contribute to the letter (visible) or to the background (not visible). As a result of the 3-frame lifetime, the number of simultaneously visible checks increased from 25 on the first frame, to 50 on the second, and to 75 on the third, after which it remained constant. Because each check appeared only once, the effective completeness at any single frame was approximately 0.12× (or 3/25^ths^ of) the designated completeness level. Consequently, in this experiment, “completeness” refers to the cumulative proportion of checks presented across the full 625 ms sequence, rather than the instantaneous completeness of any individual frame. Frames from an example dynamic stimulus are shown in Figure 10, with a sample video shown in Movie 1. Static stimuli were presented with the same 625 ms duration, with the incomplete letter stimulus shown all at once and unchanged throughout. Stimuli were presented at one of three locations: the fovea, or at 10° or 20° eccentricity in the upper visual field.

**Figure 10.**
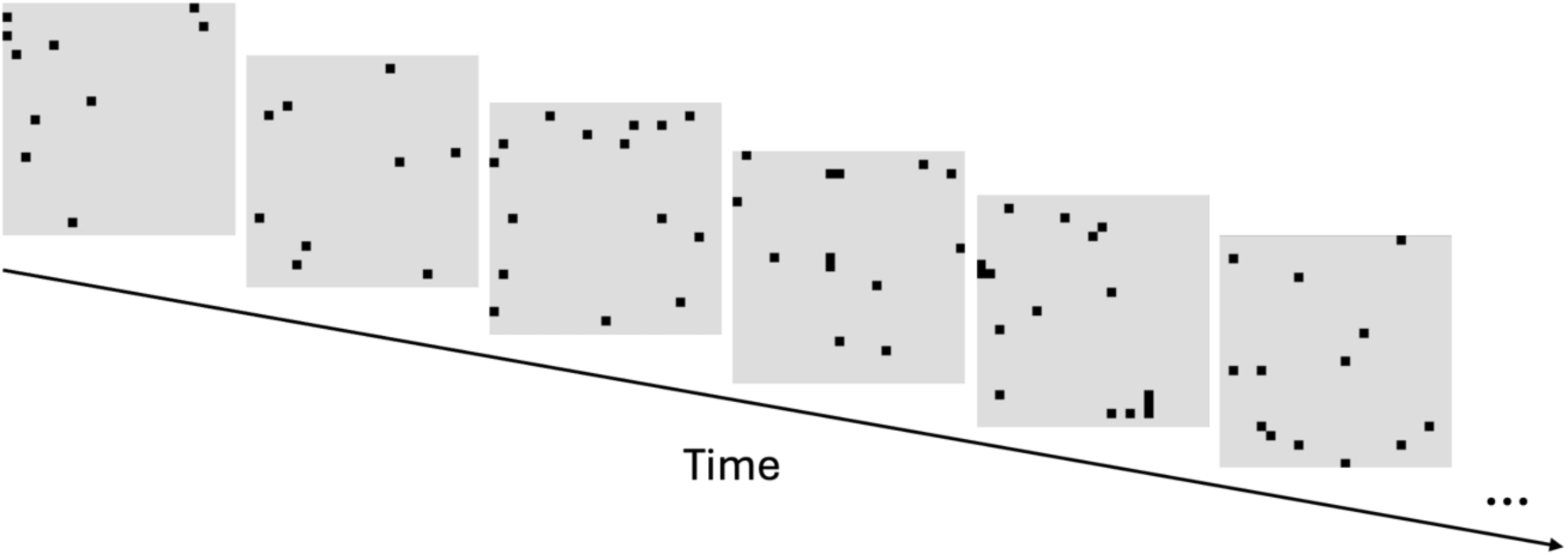
Depiction of the frames in an example trial in Experiment 3 with dynamic incomplete letter stimuli. Here, the letter “E” is shown at 0.8 completeness (when summed across the whole stimulus presentation). A new subset of checks is revealed on each frame, with checks shown for a 3-frame lifetime.

**Movie 1.**
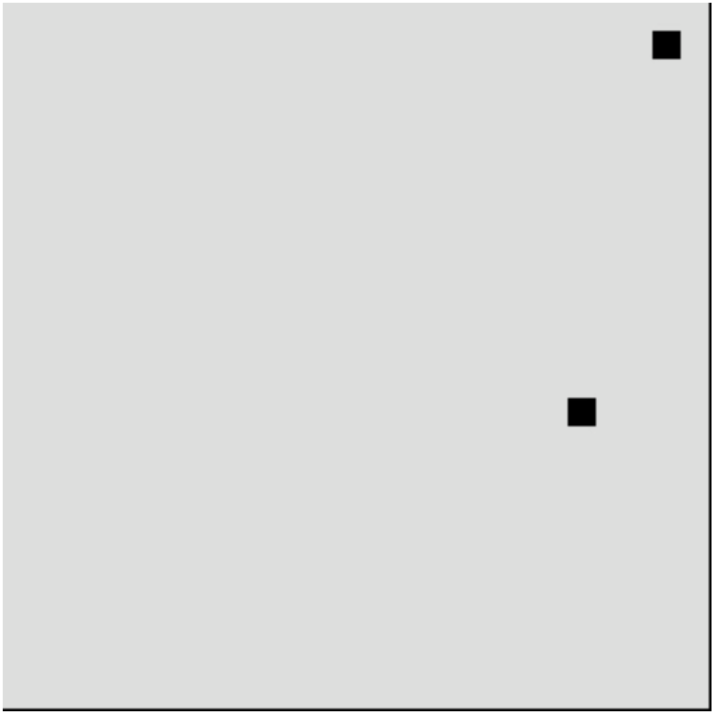
An example trial from Experiment 3 with dynamic incomplete letter stimuli. Here, the letter “E” is shown at 0.25 completeness.

Eye tracking was used to ensure fixation during peripheral presentation. The experiment consisted of six conditions (2 presentation modes × 3 eccentricities), each repeated three times. Presentation mode (static/dynamic) and location were held constant within each block. Each block contained 55 trials, including 5 practice trials at the start and 5 catch trials distributed within the block, as before running QUEST to reach threshold at 54.2% correct. The full experiment took approximately 2.5 hours, split across two sessions.

##### Data analysis

Data were analysed in MATLAB. Completeness thresholds were examined following the same procedures as in Experiments 1 and 2. Baseline performance for participant ZZ (static stimuli in the fovea) for the initial block deviated considerably from performance in the other two blocks, suggesting practice effects. This initial block was excluded from the analysis. Statistical analyses were performed using a three-way mixed-effects analysis of variance (ANOVA), with fixed effects of presentation mode (static, dynamic), eccentricity (0°, 10°, and 20°) and their two-way interaction, and a random effect of participants.

#### Results

Completeness thresholds are shown in Figure 11. As before, thresholds systematically increased with eccentricity, with poorer performance in peripheral vision compared to the fovea. Additionally, performance declined markedly (i.e. thresholds rose) with the dynamic presentation compared to the static presentation. When these factors were combined, a clear interaction was observed: the disruption from dynamic presentation increased as eccentricity increased. The ANOVA confirmed these effects, revealing significant main effects of eccentricity (*F*(2,35) = 11.26, *p* < 0.001) and presentation mode (*F*(1,35) = 21.18, *p* < 0.001), as well as a significant interaction between the two factors (*F*(2,35) = 7.85, *p* < 0.001). The main effect of participant was not significant (*F*(5,35) = 1.48, *p* = 0.323), and there was no significant interaction between eccentricity and participant (*F*(10,35) = 1.55, *p* = 0.250), suggesting consistent effects of eccentricity across participants. However, a significant interaction between presentation mode and participant (*F*(5,35) = 4.08, *p* = 0.028) indicated individual variability in the dynamic condition. In particular, two participants performed slightly worse than others in the periphery under dynamic presentation, though their performance with static presentation was comparable to the group. Baseline thresholds with static presentation in the fovea averaged 0.033 completeness, similar to the baseline performance in Experiment 2 and again lower than that reported in Yong et al. (2024). In the most challenging condition (dynamic presentation at 20° eccentricity), thresholds increased to an average of 0.4 completeness.

**Figure 11.**
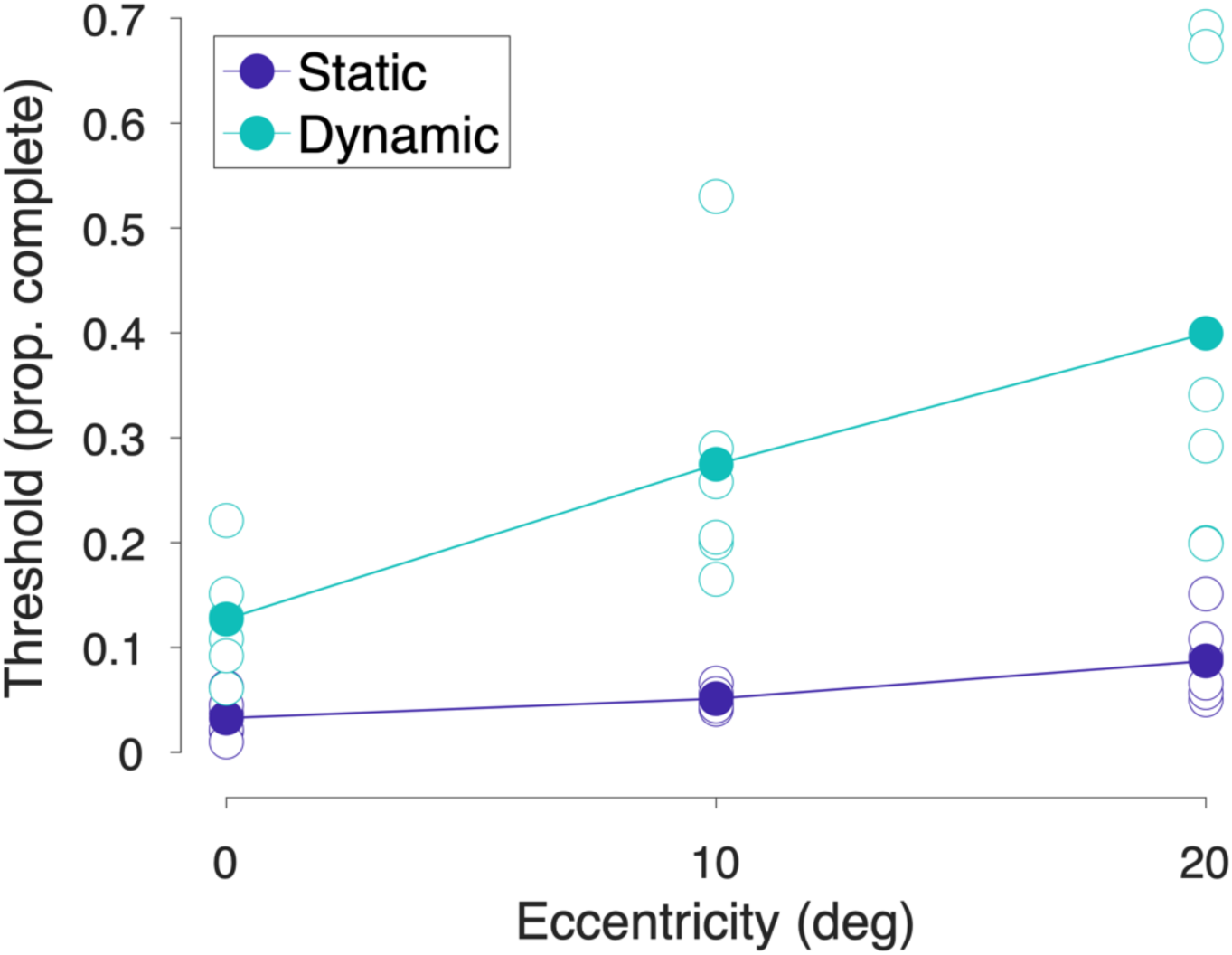
Completeness thresholds for static (blue) or dynamic (teal) incomplete letters at different eccentricities in Experiment 3. Solid dots show the mean threshold for each condition; hollow dots show thresholds for each participant.

Overall, the findings from Experiment 3 align with those from Experiment 2. Namely, both increased crowding (via peripheral presentation) and difficulties in global integration (via temporally distributed stimuli) significantly impaired incomplete letter recognition, particularly through their interaction. In this case, completeness thresholds increased up to 0.4 in the most challenging condition (dynamic presentation at 20° eccentricity), closely approximating the impairments observed in PCA patients (median: 0.47). Notably, comparable deficits in recognising incomplete letters have previously been linked to parietal cortical lesions across a range of aetiologies (Warrington & James, 1967), underscoring the cortical basis of these impairments. These results provide further support for the hypothesis that poor recognition of incomplete letters is driven by cortical factors.

## Discussion

Our aim was to investigate whether the recognition of incomplete letter elements is limited primarily by optical or cortical factors, which we approached by adapting the Graded Incomplete Letters Test (Yong et al., 2024) to selectively impose these limitations in typical adults. In Experiment 1, we observed that blur and low contrast have very little effect on GILT thresholds, producing only minor elevations when stimuli are almost at the limits of visibility. These findings suggest that optical factors are unlikely to account for the poor incomplete letter recognition observed in neurodegenerative syndromes such as Posterior Cortical Atrophy, Dementia with Lewy bodies, and Corticobasal Syndrome. The results of Experiments 2 and 3 demonstrate that cortical factors such as increased crowding and impaired global integration produced far stronger threshold elevations, to the extent that the performance of typical adults could be matched with the deficits observed in PCA. We suggest that these cortical factors play a critical role in limiting incomplete letter recognition, particularly when both factors occur together.

Our findings indicate that simple visibility issues (such as increased blur or reduced contrast) are not the primary factors limiting performance in incomplete letter recognition. In Experiment 1, blur and contrast manipulations had little-to-no effect on incomplete letter recognition, with minimal impairments (up to 0.08 proportion completeness) arising only when stimuli were presented at blur/contrast values near to thresholds for the detection of complete letters (i.e. where stimuli approached visibility limits). Incomplete letter recognition returned to baseline levels with even small increases in contrast or decreases in blur, suggesting that substantial optical degradation would be required before performance was noticeably affected. Indeed, the levels of blur and contrast reduction required to produce performance decrements exceeded those typically observed in patients with glaucoma (Crabb et al., 2013; Stamper, 1984) or cataracts (Shandiz et al., 2011). The large stimulus size used in the GILT makes it further unlikely that the central visual losses typically found in cataract and glaucoma would yield selective impairments on this task (Klein et al., 1995). This is consistent with prior findings by Yong et al. (2024), who reported that glaucoma and cataract patients performed similarly on the GILT to typical adults in the UK Biobank, despite having reduced visual acuity and contrast sensitivity. These results, together with our own, support the conclusion that optical factors are not a major limitation on GILT performance. Taken further, common age-related comorbidities in conditions like glaucoma and cataract are unlikely to be mistaken for the deficits picked up by the GILT, making it suitable to distinguish PCA and other visual-led dementias from these conditions.

Conversely, cortical factors such as crowding and global integration appear to play a critical role in recognizing incomplete letters. In Experiments 2 and 3, performance varied markedly when these factors were manipulated, far more than in Experiment 1. Both elevations in crowding and impairments in global integration impaired incomplete letter recognition, particularly when the two factors were applied simultaneously. In both Experiments 2 and 3, recognition performance worsened as stimuli were presented further from the fovea. This drop in performance is consistent with well-established eccentricity-dependent increases in crowding (Bouma, 1970), given that crowding presents the major limitation on visual perception in peripheral vision (Rosenholtz, 2016). This peripheral crowding likely disrupted performance on the GILT by interfering with the recognition of checks and their spatial arrangement/texture, thereby obscuring critical features of the letters. These disruptions are similar to the self-crowding effects seen with other stimuli (Martelli et al., 2005; Zhang et al., 2009), suggesting that this disruption may also occur within incomplete letters. We therefore suggest that crowding is a key contributor to the difficulty of recognizing incomplete letters, and in turn that the elevated crowding observed in PCA patients (Crutch & Warrington, 2007; Crutch et al., 2016; Strappini et al., 2017; Yong, Shakespeare, Cash, Henley, Nicholas, et al., 2014) may drive at least part of their impairments in GILT performance.

In addition to crowding, impairments in global integration also reduced GILT performance. Experiments 2 and 3 demonstrated these impairments using two methods that challenged global integration spatially and temporally, respectively. In Experiment 2, global integration was manipulated by altering check size. Relative to the standard check size (5 checks/stroke), performance improved with smaller checks and deteriorated with larger checks. Deficits with larger check sizes were driven by the concentration of pixels into bigger clusters, removing key features of the letters (such as corners, strokes, and gaps) more rapidly as completeness decreased. In contrast, smaller check sizes distribute the degradation across the letterform, allowing sections of these features to remain visible across a wider range of completeness levels and reducing the challenge to global integration. In Experiment 3, dynamic presentation was used to further challenge global integration. Relative to the static presentation of incomplete letters, dynamically revealing the check components of incomplete letters over time led to substantial performance impairments, even in the fovea (where crowding has little effect), demonstrating that impairments in global integration alone can challenge incomplete letter recognition. This finding aligns with the results of Zhang et al. (2024), who showed that the sequential masking of local features strongly impaired the integration required to recognise fragmented letters. In PCA, these impairments to global integration may result from disrupted recurrent hierarchical processing within the visual system or the impaired accumulation of perceptual evidence over space and time, consistent with prior proposals (Metzler-Baddeley et al., 2010; Uhlhaas et al., 2008). These spatial and temporal manipulations also offer potential strategies both to improve incomplete letter test performance in patients with PCA by supporting global integration (where manipulations maintain the spatial distribution of visual information instead of degrading local features) or to make it more difficult (potentially increasing the specificity and sensitivity of tests for these deficits).

The disruptive effect of impaired integration increased with eccentricity with both spatial and temporal manipulations, suggesting that crowding and global integration may interact rather than making independent contributions. Notably, participants faced the greatest difficulties in Experiment 3 with dynamic presentation at the farthest eccentricity, to the extent that thresholds aligned closely with the median performance of PCA patients reported by Yong et al. (2024). By pairing these deficits, we can therefore make the visual abilities of healthy 20-year-old individuals resemble those of individuals with PCA. Taken together, we suggest that deficits in incomplete letter recognition derive from processes at multiple levels: low-to-mid level processes such as crowding and higher-level processes such as global integration. As above, this is consistent with prior observations that people with PCA experience both elevated crowding (Crutch & Warrington, 2007; Crutch et al., 2016; Strappini et al., 2017; Yong, Shakespeare, Cash, Henley, Nicholas, et al., 2014) and difficulties integrating visual features across space and time (Metzler-Baddeley et al., 2010; Uhlhaas et al., 2008). Our findings suggest that both of these factors may be insufficient to explain the full magnitude of GILT performance deficits on their own, with patient performance only approached when the two are combined. It may be then that deficits at multiple levels of the visual system contribute to the difficulties PCA patients face in recognizing incomplete letters.

It is nonetheless possible that impairments in incomplete letter recognition arise from other visual processes than those discussed above. In particular, low-to-mid level processes such as contour integration (Field et al., 1993) and subjective contour recognition (Kanizsa, 1976) may play a role in influencing the recognition of incomplete visual stimuli. Contour integration refers to the ability to link or bind visual elements together to form coherent perceptual groupings, a process that requires the integration of information across neighbouring filters tuned to similar orientations (Field et al., 1993). PCA patients exhibit significantly higher thresholds for contour integration than typical participants, indicating deficits in this process (Metzler-Baddeley et al., 2010). However, such deficits may reflect more general impairments in visual integration, including spatial feature binding and/or the formation of global shape representations, which would also likely disrupt incomplete letter recognition, as argued above. Crowding may similarly interfere with contour-integration processes by disrupting the spatial and feature-level coherence needed for perceptual grouping (Chakravarthi & Pelli, 2011; May & Hess, 2007). It is also possible that alternative higher-level deficits observed in PCA could contribute to impaired incomplete letter recognition. For example, simultanagnosia (difficulty perceiving multiple objects at once; Coslett & Saffran, 1991; Primativo & Starrfelt, 2025) and metamorphopsia (localised distortions to objects in the visual field; Crutch et al., 2011) could impair performance in incomplete letter recognition. However, as with contour integration, these higher-level deficits may also be driven by elevations in crowding (Strappini et al., 2017), making the precise direction of their involvement difficult to ascertain. Nonetheless, our findings suggest that both low- to mid-level processes (such as crowding) and higher-level processes (such as global integration) are strong candidate mechanisms underlying the deficits in PCA patients. Finally, although higher-level cortical functions such as working memory and executive function can be affected in PCA (Crutch et al., 2017), a strength of the GILT is that the simple and intuitive letter-identification task should minimise demands on these processes. Any contribution of higher-order cognitive factors would therefore be expected to disrupt the recognition of both complete and incomplete letters, i.e. manifesting as errors even at 100% completeness (Cummings et al., 1986; Yong et al., 2016). Accordingly, participants with AD show typical threshold levels and clear task comprehension regarding the required letter response, despite difficulties with memory and executive function (Yong et al., 2024). These issues are especially unlikely to explain the present findings in typical adults, particularly because cognitive demands were constant across conditions.

In conclusion, our findings demonstrate that optical factors have little-to-no limiting effect on impaired performance in incomplete letter recognition. This outcome aligns with the observations by Yong et al. (2024), and highlights that performance impairments on the GILT task are unlikely to arise from common age-related optical conditions such as cataracts or glaucoma. This invariance with optical properties makes the task well-suited for detecting cortical deficits in visual processing. Beyond ruling out optical limitations, our data also support the hypothesis that poor performance in incomplete letter recognition is primarily driven by cortical deficits. We demonstrate in typical adults that elevated crowding and difficulties in global integration can strongly impair the visual system’s ability to reconstruct fragmented letter forms. At the low- to mid-level, these processes could impair both the recognition of local visual features and the perception of their spatial arrangement. At a higher level, difficulties with global integration could further limit the ability to assemble partial visual input into recognizable letter shapes. Future studies will extend this line of inquiry to include patients with neurodegenerative syndromes, with the aim of deepening our understanding of the recognition deficits in this population.

